# Reconfiguration of Behavioral Signals in the Anterior Cingulate Cortex based on Emotional State

**DOI:** 10.1101/2023.07.12.548701

**Authors:** Adrian J Lindsay, Isabella Gallello, Barak F Caracheo, Jeremy K Seamans

## Abstract

Behaviours and their execution depend on the context and emotional state in which they are performed. The contextual modulation of behavior likely relies on regions such as the anterior cingulate cortex (ACC) that multiplex information about emotional/autonomic states and behaviours. The objective of the present study was to understand how the representations of behaviors by ACC neurons become modified when performed in different emotional states. A pipeline of machine learning techniques was developed to categorize and classify complex, spontaneous behaviors from video. This pipeline, termed HUB-DT, discovered a range of statistically separable behaviors during a task in which motivationally significant outcomes were delivered in blocks of trials that created 3 unique ‘emotional contexts’. HUB-DT was capable of detecting behaviors specific to each emotional context and was able to identify and segregate the portions of a neural signal related to a behaviour and to emotional context. Overall, ∼10x as many neurons responded to behaviors in a contextually dependent versus a fixed manner, highlighting the extreme impact of emotional state on representations of behaviors that were precisely defined based on detailed analyses of limb kinematics. This type of modulation may be a key mechanism that allows the ACC to modify behavioral output based on emotional states and contextual demands.

## Introduction / Main

Performance on any task can be influenced by a number of factors that are not directly related to the task itself, including the level of hunger or thirst, stress, fatigue or frustration. This occurs because emotional states (Kragel et al 2016) and basic autonomic functions (Tu & Zhang 2022) are not isolated to particular brain regions but directly influence activity patterns throughout the brain. It is not clear exactly how this occurs at a cellular level, but a recent study by Cowley et al (2020) showed that changes in internal state variables were reflected as a ‘slow neural drift’ in the activity neurons in the primate prefrontal cortex that in turn affected how these neurons processed task-relevant information on a decision-making task. In the last 10 years that there has also been accumulating evidence that behaviors are tracked by widespread brain regions, including areas that have no obvious role in movement (Kaplan & Zimmer 2020). The reason why brain regions that do not directly control emotion, autonomic tone or actions have access to all this information it not entirely clear. However, it does suggest that the brain tends to create holistic representations of experiences rather than operating as a series of highly specialized regions, each processing information within a singular domain.

Nowhere is the integrated view of information processing more evident than the frontal cortices. Here neurons exhibit the most extreme form of mixed-selectivity (Fusi et al, 2016), as they simultaneously represent information across many domains and dynamically re-weight these representations in order to adapt to changing task demands (Rigotti et al, 2013, Barak et al., 2013, Stringer et al 2019; Kira et al 2023). This takes different forms in different subregions of the frontal cortex. Anatomical and electrophysiological data suggest that lateral areas of prefrontal cortices may prioritize cognitive and sensory streams of information whereas more medial regions such as the anterior cingulate cortex (ACC) place a greater emphasis on processing information related to emotional/autonomic states and action. This dichotomy is highlighted by the effects of localized lesions of each region as damage localized to lateral PFC, tends to produce deficits in cognition and executive functions whereas more medial lesions centered on the ACC, result in changes in emotion, autonomic control and behavioral output (Devinsky et al, 1995; Rudebeck et al, 2006; Brotis et al, 2009). Of note in this regard, surgically precise lesions of the ACC have been shown over the last 70 years to provide relief to patients suffering from diseases where an emotional or autonomic state exerts a pathological level of control over the patient’s thoughts and behaviors as in obsessive-compulsive disorder (OCD) and chronic pain conditions. Studying the multiplexing of emotional and behavioral information by ACC neurons may offer a unique window into how the brain creates holistic representations and how these representations may be altered in certain disease states.

In order to study how ACC neurons multi-lex information about emotion and behavior, we must be able to accurately identify behaviors performed in different emotional states while also being able to disambiguate the neural correlates associated with the emotional state from the neural correlates associated with the behavior. As a first step towards addressing this challenge, Lindsay et al. (2018) used various machine learning approaches to precisely quantify the motions of individual limbs from video taken above and to the side of rats in a small transparent enclosure. In that study, ACC neurons were found to encode individual limb movements and positions as well as the animal’s allocentric location, even though the animals were not required to perform any task and were simply moving around freely in the enclosure. The present study builds on that work, progressing from the quantification of the neural correlates of individual limb movements to the quantification of the neural correlates of behaviors.

In recent years, several methods have been developed to categorize temporal sequences of limb movements as distinct behaviours (Arac et al 2019; Luxem et al 2020). One factor that limits their utility for our purposes is that they typically define behaviours across a very specific temporal window. In contrast, we were interested in behaviours that evolve over variable and often unknown timescales.

Another limitation of currently available techniques is that while they all effectively parse movements into behavioural clusters (i.e. unique constellations of limb movements), it can be unclear as to what these clusters represent or mean in functional terms. Finally, many of these approaches are not designed to seamlessly relate discovered behaviours to neural activity patterns.

In response to these challenges, we developed a pipeline of machine learning techniques built on work in fruit flies (Berman et al 2014; Günel et al 2019), rodents (Hsu et al 2021), and humans (Mehta et al 2020), that allowed us to categorize or classify unique sequences of individual limb movements across variable timescales. The pipeline also enabled us to compare automatically discovered behaviours with manually selected behaviours in a shared behavioural space, which helped to facilitate the interpretability of discovered behaviours. Finally, it includes various machine learning techniques to find links between discovered behaviours and neural activity patterns.

This pipeline, termed ‘Hierarchical Unsupervised Behavioural Discovery Tool’ (HUB-DT) was used to detect the specific behaviors expressed during our ‘3-valence task’ (Caracheo et al 2018). In this task, 3 outcomes that differed in emotional valence were delivered to rats in separate blocks of trials with each block of trials involving outcomes of a single valence type. As a result, each block of trials was considered to form a unique ‘emotional context’ and was associated with a statistically separable ensemble activity-state pattern in the ACC that persisted throughout the entirety of the block, including the ∼50s inter-trial intervals where no cues or outcomes were presented (Caracheo et al 2018). HUB-DT was not only able to identify a large number of putative behaviours, but some of these behaviors were unique to a specific emotional context. It was also able disambiguate neural signals related to behaviors from the neural signals related to emotional context. Overall, we found that the majority of ACC neurons responded to actions in a contextually dependent manner. This modulation provides a potential mechanism for emotional regulation of behavioral output.

## Results

All experiments involved the 3-valence task (see Caracheo et al 2018) which was performed in a single physical enclosure (Figure 1). On each session, rats were given a tone stimulus paired with different, motivationally-specific outcome in blocks of 30 trials. Each block contained trials with a single tone/outcome pairing and was referred to as a unique ‘emotional context’. In one emotional context, a tone preceded an aversive outcome in the form of a mild foot shock (the S-context), in another emotional context, a different tone preceded an appetitive outcome in the form of food pellet delivery (the F-context), and a third emotional context block, a 3^rd^ tone preceded a neutral outcome (no outcome, the N-context). The trials were separated by an average inter-trial interval of 51s and there were no additional time gaps between contexts.

**Figure 1:**
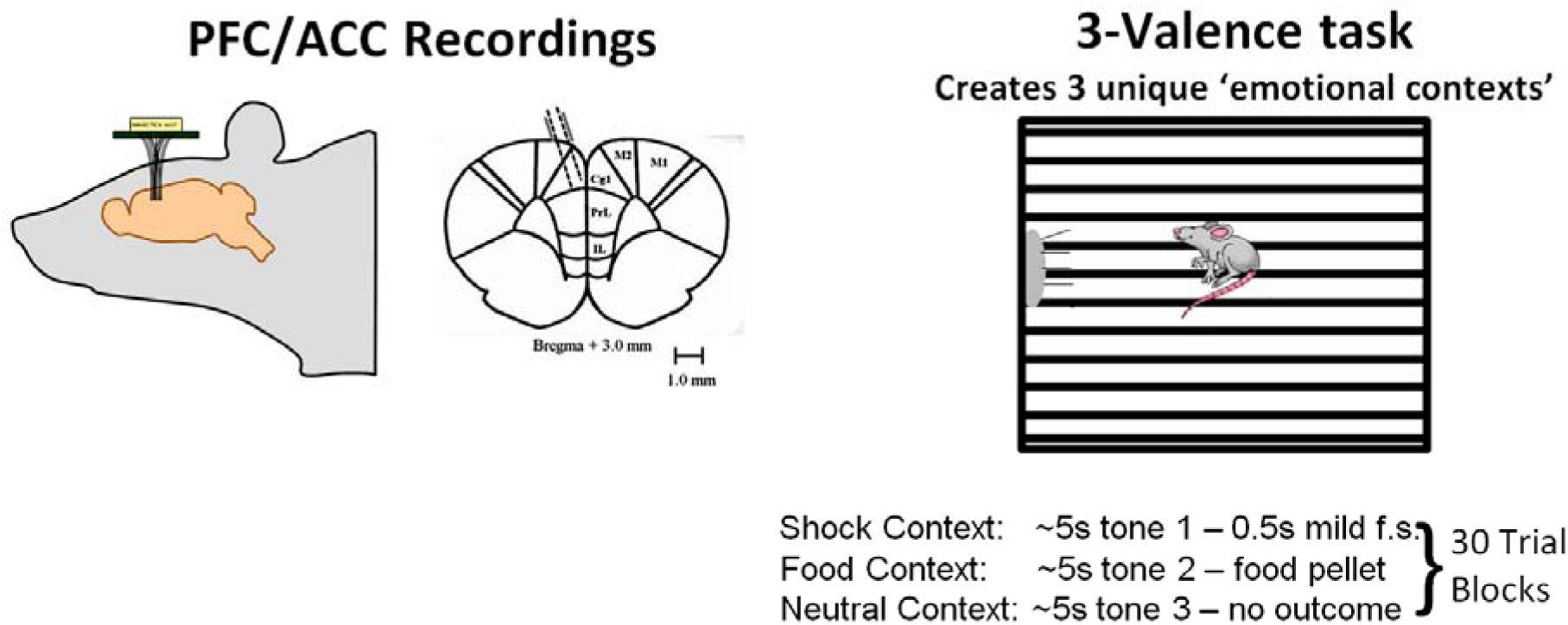
Recording sites and task schematic. **A)** Tetrodes were slowly advanced through the dorsal ACC to the level of the prelimbic cortex across sessions. **B)** A schematic of the 3-valence task. Rats were placed in a single transparent enclosure equipped with a grid floor to deliver footshocks, a speaker to deliver tones and a port to deliver food pellets. A unique tone was followed by a footshock (S-context), the delivery of a food pellet (F-context) or by no outcome (N-context). Each tone outcome pairing was delivered in a dedicated block of 30 trials that is referred to as an emotional context.

### Hierarchical Unsupervised Behavioural Discovery Tool

The Hierarchical Unsupervised Behavioural Discovery tool (HUB-DT), was used to automatically detect behaviors evolving across various timescales. Similar to motion mapper (Berman et al 2014), HUB-DT treated behaviours as a combination of features (i.e. limbs, joints, or body parts) that moved in specific patterns at particular timescales or frequencies. Specifically, HUB-DT transformed 3-d body tracking data from DeepLabCut (Mathis et al 2018) by wavelet projection with multiscale Morlets to embed each pose (i.e. the constellation of limb positions at a single point in time) into a spatiotemporal high-dimensional space, where the axes were the responses of a particular limb moving at a particular frequency (Figure 2:A-C). This space was then reduced in dimensionality using t-SNE (van der Maaten et al 2008), and points in this low-dimensional projection were clustered using Hierarchical Density-based Spatial Clustering of Applications with Noise (McInnes et al 2017: HDBSCAN). Each cluster of points in this space was considered to be a distinct, putative behaviour. To give a concrete example, if there were multiple occasions where the right forelimb was moving at 0.2Hz, the left hindlimb at 1Hz and the head at 0.5Hz, these points would form a cluster in the high dimensional limb x wavelet space and as such would be considered a behavior. The characteristics of a given cluster could then be evaluated by generating a ‘behavioural prototype’ which was the set of average responses for the cluster (Figure 2F).

**Figure 2:**
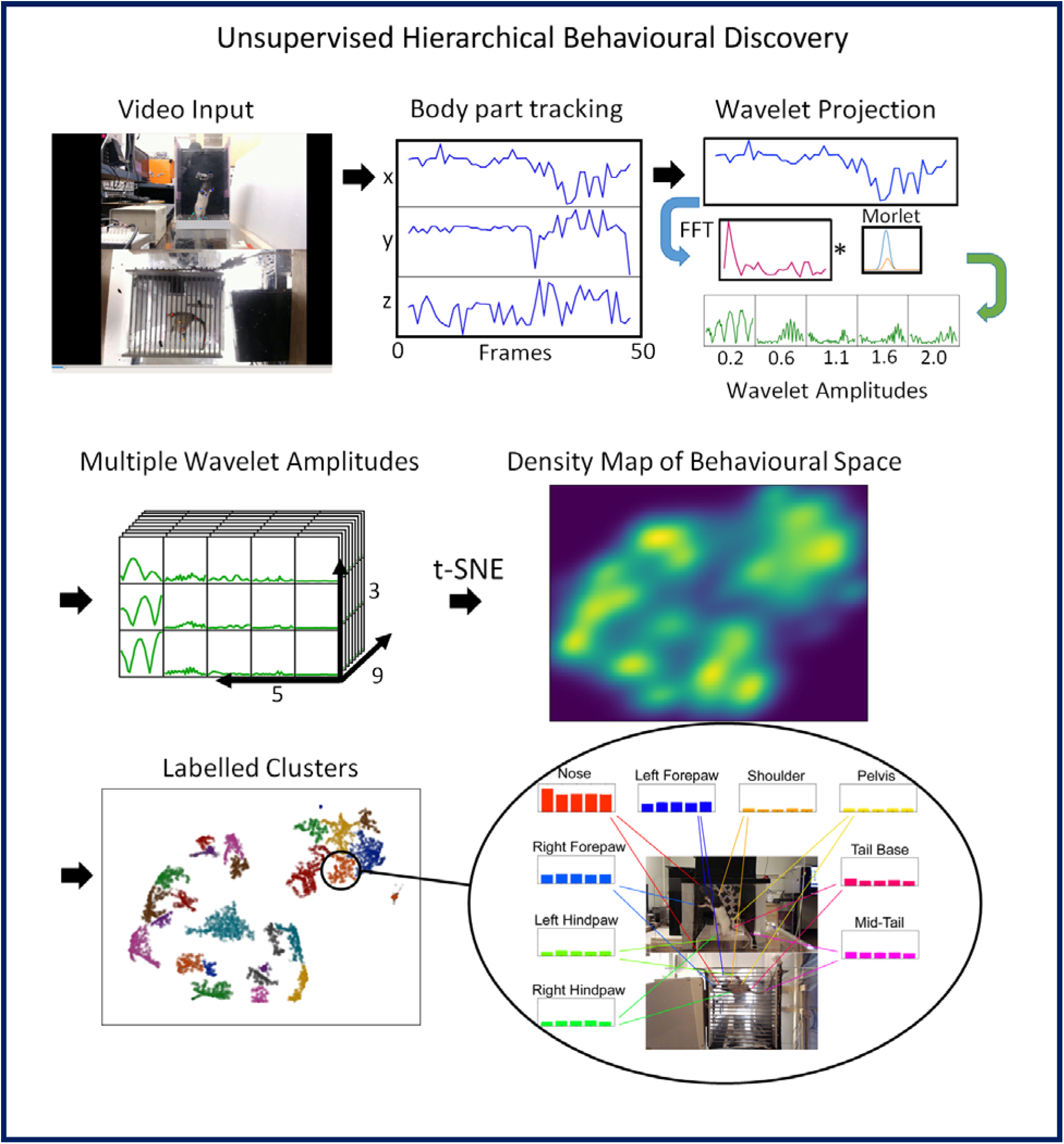
A schematic description of the HUB-DT process. **A)** Video was collected by cameras placed underneath and to one side of the transparent enclosure. **B)** (Left) The positions of 9 parts of the animal’s body were tracked using DeepLabCut (all positions were normalized relative to the rat’s center of mass). (Right) The x,y,z coordinates of each body part were then convolved with 5 logarithmically-distributed frequency Morlets (ranging from 5Hz to 0.5Hz). **C)** This resulted in a behavioural space of 135 dimensions (‘9’ body parts x ‘3 x,y,z’ coordinates x ‘5’ Morlets). **D)** The 135-dimensional space was projected down to 2 dimensions using t-SNE to produce a density map. **E)** HDBSCAN was applied to the density map to provide unsupervised clustering: each cluster represents a distinct behavior. **F)** Each cluster can be characterized by a prototypical or average set of wavelet responses corresponding to the multi-timescale movements of body parts in the putative behaviour. In this example, the behavior was characterized by movement of the nose, right and left forepaws and base of the tail across multiple frequencies with little involvement from the other body parts. Although we often prefer to refrain from assigning a human label for the discovered behaviors, this profile is roughly consistent with what would typically be considered exploration/rearing.

### Discovering Behaviour at Variable Granularities

A feature of hierarchical density-based clustering is its ability to assign cluster labels at variable densities. Within HUB-DT, clusters were created within the behavioural space based on the relative distance of points to other points in the local neighborhood, specified as a ‘reachability distance’. This distance relationship was then expressed within a reachability tree, that described the hierarchy of clusters. The tree could be ‘sliced’ (i.e. extraction of cluster labels at a fixed depth in the tree) at various levels, which would be functionally equivalent to the fixed density clustering implemented by the standard DBSCAN algorithm. This made it possible to generate clusters at specific neighbourhood density thresholds, in-effect allowing us to control the granularity of the discovered behaviours (Figure 4). Shallower slices (higher up the tree) resulted in clusters with reduced granularity, as characteristics separating behaviours further down the tree would be merged together. As the fixed-density line descends, more and more clusters emerge, separated by increasingly subtle differences in limb kinematics. At full depth, the algorithm selects clusters adaptively, based on their persistence and relative density. Unless otherwise stated, all analyses below involved behaviors detected at full depth of the hierarchy.

**Figure 3:**
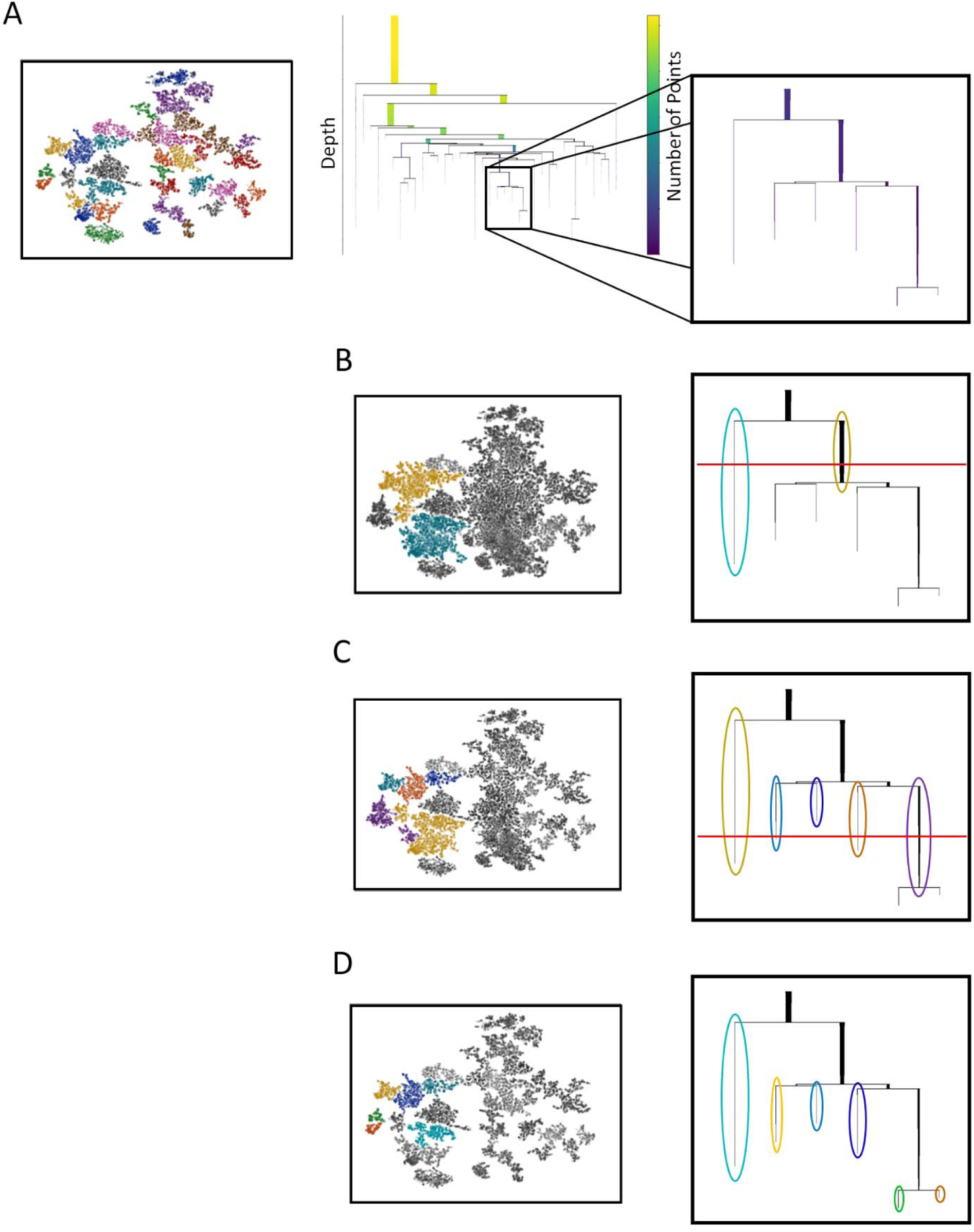
Behavioral discovery at different levels of the behavioral cluster hierarchy. **A)** (Left) A set of clusters (labelled by colour) generated by HUB-DT plotted over the 2-D density map. (Mid) Behavioral cluster hierarchy as a dendrogram, with the width and color of each branch representing the number of points in the cluster at that level. Unlike a standard hierarchical tree where a cluster splits into two new clusters at each level, HDBSCAN generates a condensed reachability tree tracking persistent clusters that are continuously shedding points while moving downward along the density gradient. This is depicted by the fading/narrowing of each line. The box (Right) draws attention to one segment of the tree. **B-D** We follow this segment and the associated clusters through progressively deeper slices of the tree. (Left) The clusters with this specific segment. (Right) The tree segment showing a line of fixed density (red) that can be drawn through the tree to extract cluster labels at a certain level. Shallow (B), or deeper (C), or at full depth (D) increasing the granularity of the labelled behaviours. Cluster separation and membership is based on neighbourhoods in the density map result from differences in limb kinematics.

**Figure 4:**
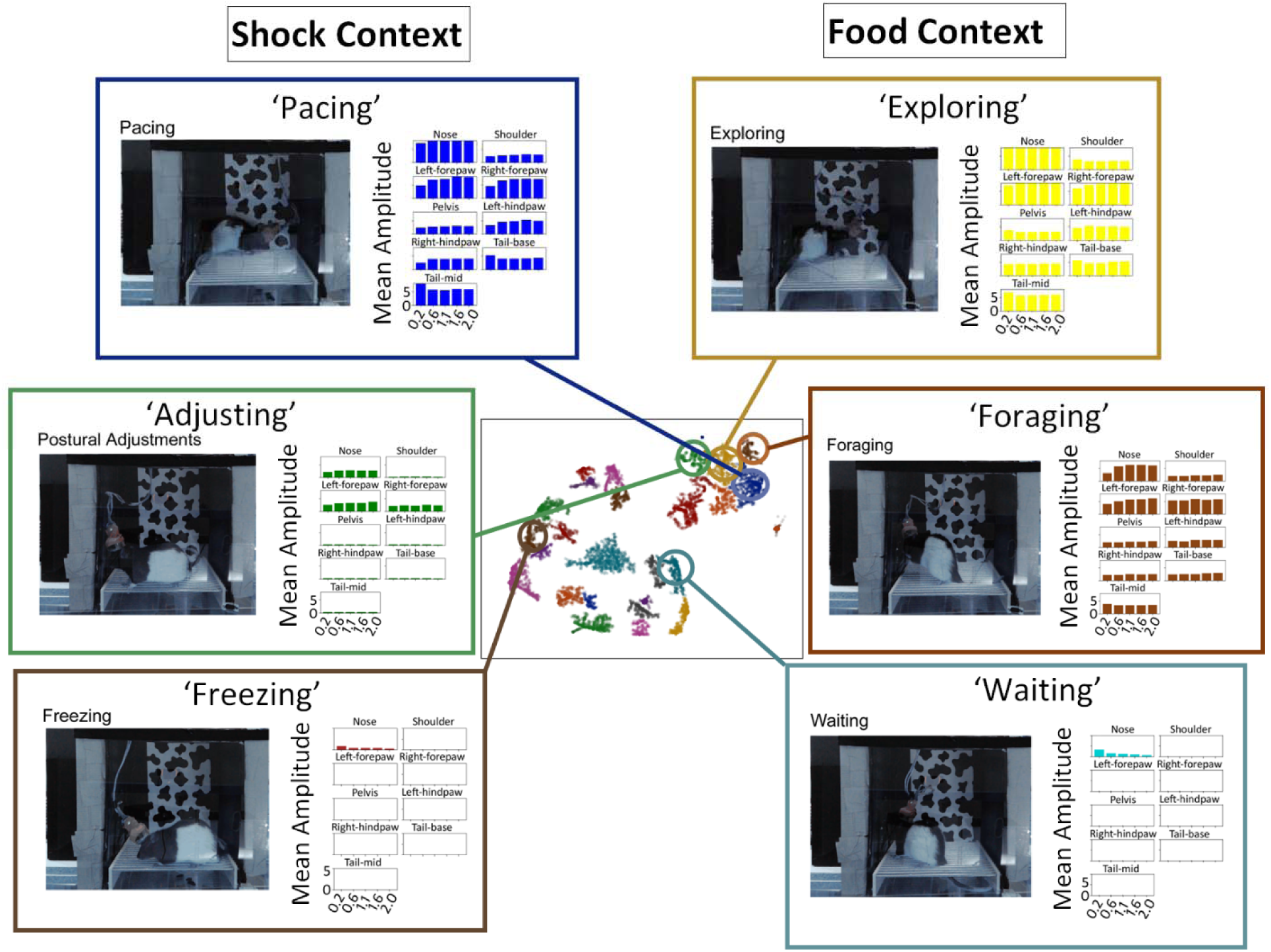
Identifying different behaviors detected in a single session. The inner subpanel shows the clusters of behaviors produced by HUB-DT and the outer subpanels show 6 of these behaviors in greater detail. Each of these behaviors has been assigned an interpretable name based on extracted video snippets. The image on the left inset is a screenshot from an example video when the animal was engaged in the behavior while the right inset panel gives the associated behavioral prototype: The behavioral prototype is the mean of the cluster responses, and gives the average amplitude across the 5 wavelets for each of the body parts. The 3 behaviors on the left were specific to the S-context and the 3 behaviors on the right, to the F-context, but this information was not used by HUB-DT to differentiate them.

### Identifying Discovered Behaviours

The behaviors detected by HUB-DT did not always fit into categories that could be assigned familiar names. It was nevertheless still possible to understand the detected behaviors based on their defining features. Figure 3 shows 6 behaviors identified by HUB-DT. In each panel, the behavioral prototypes (i.e. the average of the behavioral cluster) are shown on the right. Note that these prototypes differed both within and across emotional contexts. In particular, ‘pacing’, ‘exploring’ and ‘foraging’ all involved locomotion but were associated with slight differences in limb kinematics. Likewise, ‘freezing’ and ‘waiting’ appeared to be virtually identical but were differentiated by HUB-DT based mainly on subtle differences in movements of the nose/snout. It is also interesting that these two behaviors were performed in only either the S-context or F-context respectfully, even though this information was not used by HUB-DT to differentiate them. Conversely, there were multiple behaviours that were repeated across the 3 contexts, as described in more detail below.

### Comparing Human and Automatically Generated Behavioural Labels

Two analyses were used to compare the performance of HUB-DT to that of a human observer scoring behaviours from video. The first focused on the assumption that each emotional context should evoke at least some behaviors that were expressed preferentially in that context. The emotional context specificity of HUB-DT versus human-labelled behaviours was assessed by a two-way cluster purity measure which quantified the proportion of a given behavioral cluster that was contained within a particular emotional context cluster versus the proportion of the emotional context cluster that was accounted for by the behavioral cluster. We found a substantial advantage in peak specificity for the HUB-DT discovered behaviours (81%±4.7) as compared to manual behaviour labels (43%±9.2; t=142.4, df=8, p<0.05) (Figure 5). This meant that HUB-DT detected behaviors that were more context-specific than typical pre-selected behaviours scored by human observers.

**Figure 5:**
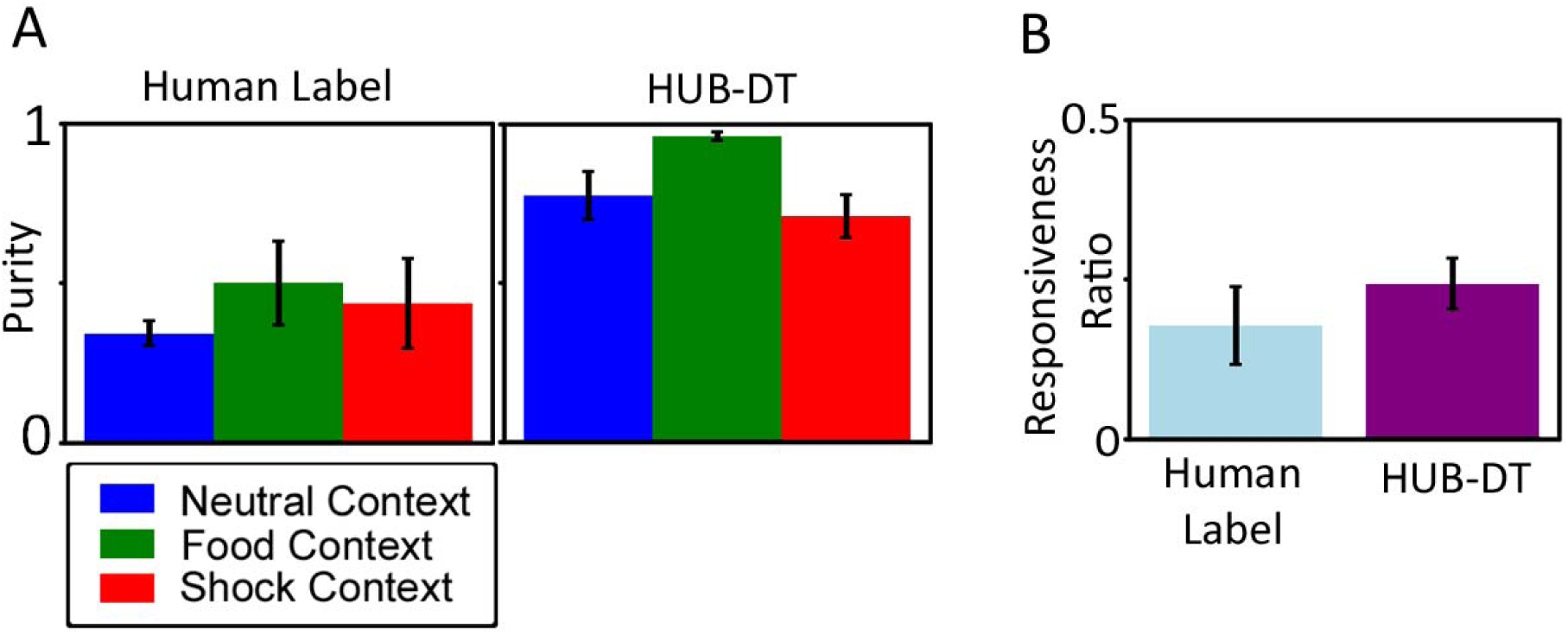
Comparing behavioral labels assigned by a human versus HUB-DT. **A)** A two-way purity measure showing the context specificity of behaviors. On the left is peak purity for pre-selected human annotated behaviours, averaged across sessions. On the right, peak purity for discovered behaviours averaged across all sessions. **B)** GLMs were used to detect neurons that were significantly responsive to a given behavior. The responsiveness ratio which compared the number of GLM-responsive versus GLM-nonresponsive neurons did not differ significantly for behaviors detected by HUB-DT versus a human observer

The second comparison assessed the responsiveness of single neurons to a given set of labels, if the labels were generated by a human observer versus HUB-DT. This was assessed by multifactor Generalized Linear Models (GLMs) as follows. All time bins within a session had a human-generated behavioral label and a HUB-DT generated behavioural label, along with a set of associated firing rates. Since a human did not manually annotate all sessions, we first propagated their behavioural labels across all datasets using a Random Forest (RF) classifier (see Methods). The GLMs were then trained to relate single neuron firing to human-generated or HUB-DT generated behavioural labels. The strength of these relationships were quantified using the coefficient of determination (r^2^) and decisions about whether a given neurons was responsive to a given factor were based on threshold criteria (see Methods). This allowed us to compare the ‘responsiveness ratio’ (the ratio of GLM-responsive to GLM-nonresponsive neurons) across the two sets of labels. While the HUB-DT discovered behaviours had a higher overall mean responsiveness ratio (24%±3) than the human behavioural labels (17.8%±5.4), this difference was not significant (F(df)=9, p=0.35; Figure 5). Based on these comparisons, we suggest that HUB-DT can discover behaviours that are significantly more context-specific and slightly more representative of ACC firing than human-labelled behaviours. The remainder of the study will focus only on behaviors detected by HUB-DT.

### Disentangling Behaviour and Emotional Context Signals in Single Neurons

A similar GLM-based approach was used to determine whether individual neurons were responsive to behaviors, to emotional contexts or some combination of the two. Time bins in each session were assigned three sets of labels, one for the overall emotional context (S-context, F-context, N-context), one for the task-event (S-tone, F-tone, N-tone), and one for the HUB-DT assigned behavioural label (Figure 6). A series of multifactor GLMs were then trained to relate single neuron firing to one of the three sets of labels. Multiple GLMs, one for each factor set, were used in order to avoid multicollinearities caused by the non-independence within and across time bins. Single neuron firing rates were predicted by these GLMs, yielding a measure of responsiveness of a neuron to a given factor where responsiveness was the proportional contribution of the neurons’ firing rate to a given GLM factor (with a threshold to address numerical instability and to remove low-likelihood correlations; see Methods). These analyses involved a total of 263 neurons recorded across 5 sessions. Overall, 51.7% of neurons were responsive to an emotional context contrast, with 12.1% being responsive to emotional context contrasts isolated to the tone epochs (Figure 6). In addition, 39.1% of neurons were responsive to at least one of the behaviors. However, it is important to emphasize that most neurons were responsive to one or more contexts (inside or outside of the tone epoch), one or more behaviors or some mixture of the two categories (Figure 6). In fact, only 28.9% of neurons were responsive to a single emotional context (including the tone epoch) and only 2% to a single behavior. On the other hand, of the 142 neurons responsive to emotional contexts (and tones), 64 (45%) were also responsive to a behavior. This highlights the multi-modal nature of ACC representations, both in terms of inter-mixing different emotional valences as well as inter-mixing emotional and behavioral information. Additionally, although a large proportion of neurons were responsive to at least one of the categories, the magnitude of the association to a given factor (i.e. the r^2^) was quite small for each neuron, as on average, 4.9 ± 1.5% of a neuron’s firing was related to the encoding of behaviours and 7.5 ±1 % to emotional context.

**Figure 6:**
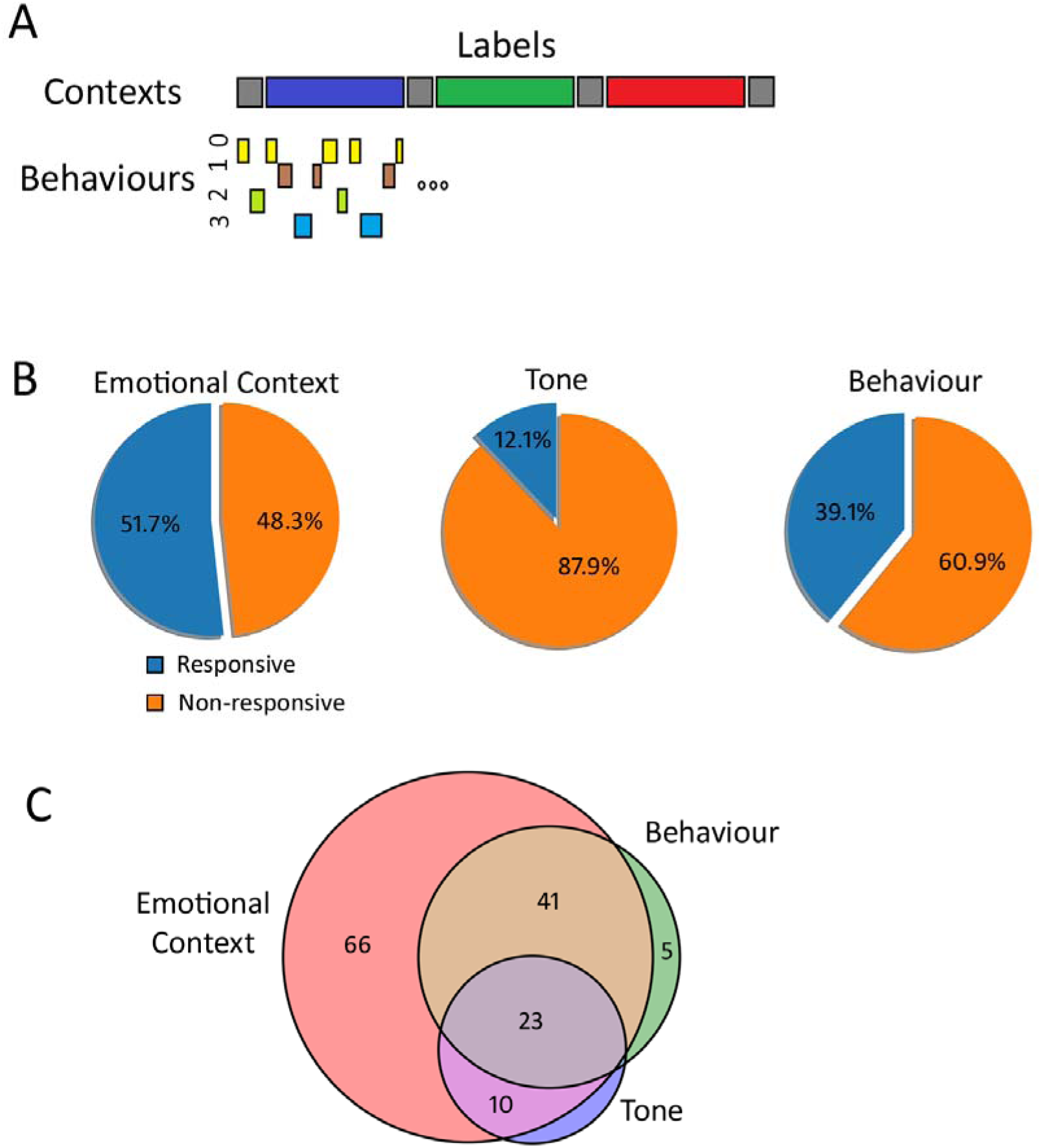
Measuring Responsive Neurons across the population. (A) A schematic showing the factors for Generalized Linear Model (GLM) analysis. Each time bin has an associated label for emotional context, behaviour (from HUB-DT), and task factors. (B) Proportions of responsive and non-responsive (as measured by the coefficient of determination from a trained GLM) neurons across the population for three sets of factors: Emotional Context, Behaviour, and Tone. (C) Responsive neurons plotted as a Venn diagram highlighting the distribution of multi-selectivity in the population. While the bulk of responsive neurons were responsive to emotional context, there was significant overlap with other factors; nearly half of emotional context responsive neurons were also responsive to a behaviour.

It is possible that some of the emotional context responses may have arisen because the neurons were tracking behaviors that were performed in only a single emotional context (See figure 3). In order to address this potential confound, we again used a GLM approach, but in this case trained a second GLM to predict emotional context labels using the residuals from the GLM above that used behavioral labels as factors. In other words, this second GLM was asked to predict emotional context, after the contribution to behavioural encoding had been removed. The strength of emotional context encoding was quantified using the coefficient of determination (r^2^). Results showed that there was no change in contextual responsivity for the GLM trained on the behavioral residuals versus a GLM trained on raw firing rates (Figure 7; t=0.583, df=257, p>0.05). The fact that the results from these two GLM analyses did not differ suggests that even though ACC neurons could be responsive to behaviors in a context-specific manner, this was not the information these neurons used to differentiate the emotional contexts.

**Figure 7:**
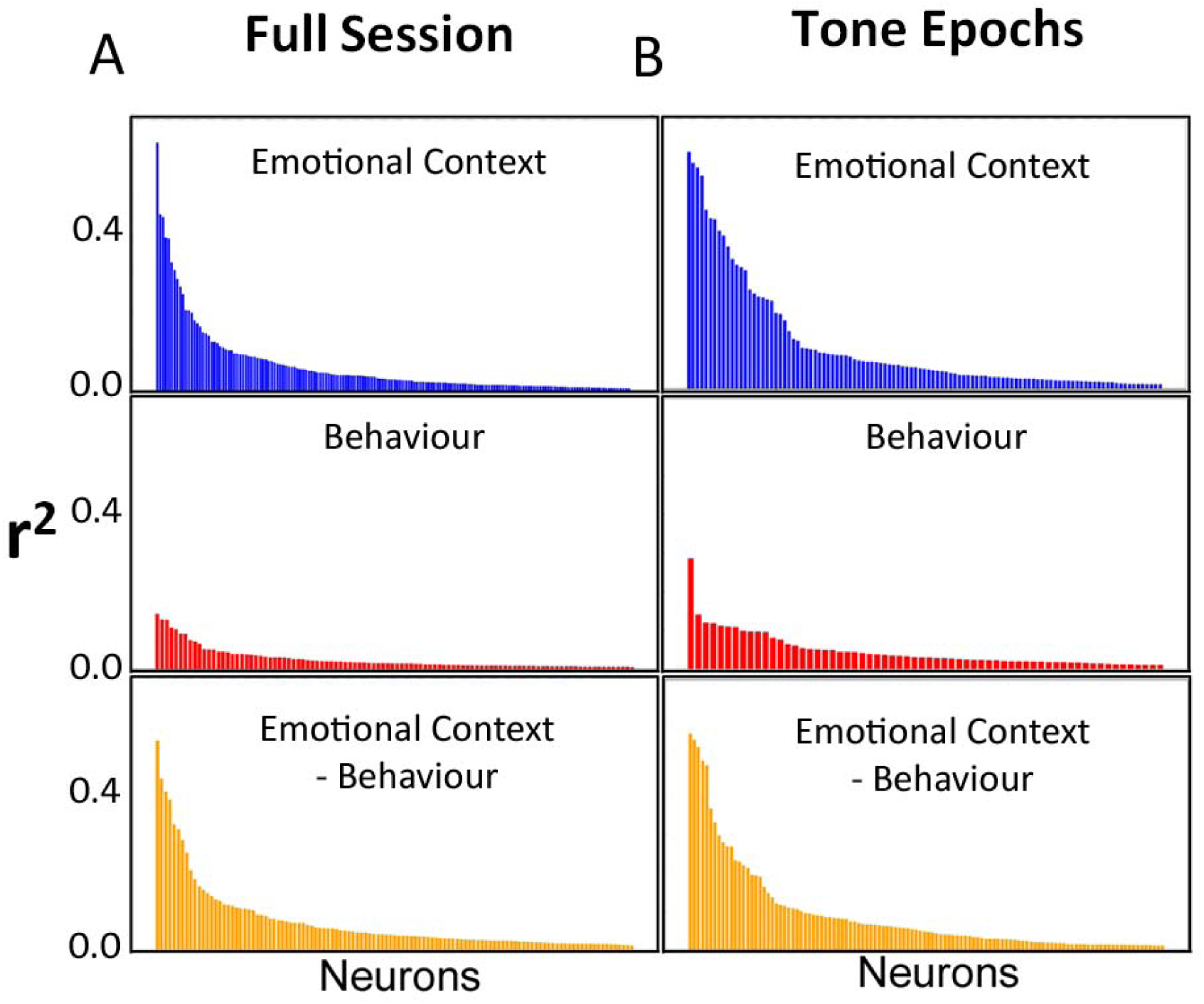
The impact of context-specific behaviors on contextual encoding by individual ACC neurons. Multifactor GLMs were trained to relate single neuron firing to a set of labels contrasting the 3 emotional contexts, and to a set of labels for HUB-DT behaviours. The labels denoted every time bin within full sessions (A) or the time bins associated with just the conditioned tone epochs (B). Each panel shows sorted r^2^ values that relate spike counts to the factors for all neurons. The Top panels (Blue) give the r^2^ values when the GLM was trained using raw spike counts on emotional context. The middle panels (Red) when the GLM was trained using raw spike counts on behaviours. The Bottom panels (Orange) give the r^2^ values when the GLM was performed using spike counts in which the contribution to all behavioral factors had been removed (i.e. the residual spike count matrix from the GLM trained using the detected behaviors as factors).

### Predictive Decoding from Populations of Neurons

As a follow up to the single neuron analyses, we investigated representations of behaviors and contexts at the ensemble level using a predictive decoding framework. Specifically, Random Forest (RF) models were trained to predict behavioural and emotional context labels from ensemble firing using all neurons in a given session. Using these RFs, we found that ensemble firing was able to decode both behaviours (93%±0.5 accuracy averaged across behaviours and sessions) and emotional contexts (82 ± 1.6% accuracy averaged across sessions and contexts) in unseen samples with an accuracy much higher than for any single neuron in isolation (Figure 8). Similar to the single-neuron analyses above, we addressed the question of whether context-specific behaviors contributed to the accuracy of context-block separations by testing whether the decoding of emotional context changed when it was performed using raw firing rates versus rates where the contribution of behavioral factors had been removed (i.e. using the residual firing from single-neuron behavioral label GLM described above). When the residuals from the behavioral label GLM were passed to the RF models, a mean emotional context decoding of 85% ± 1.2 was attained across sessions, as compared to 82% ± 1.6 when the raw firing rates were used (Figure 8). This modest improvement in performance again indicates that ACC ensembles did not differentiate emotional contexts by tracking context-specific behaviors. In fact, in this case behavioral encoding slightly hampered emotional context differentiation.

**Figure 8:**
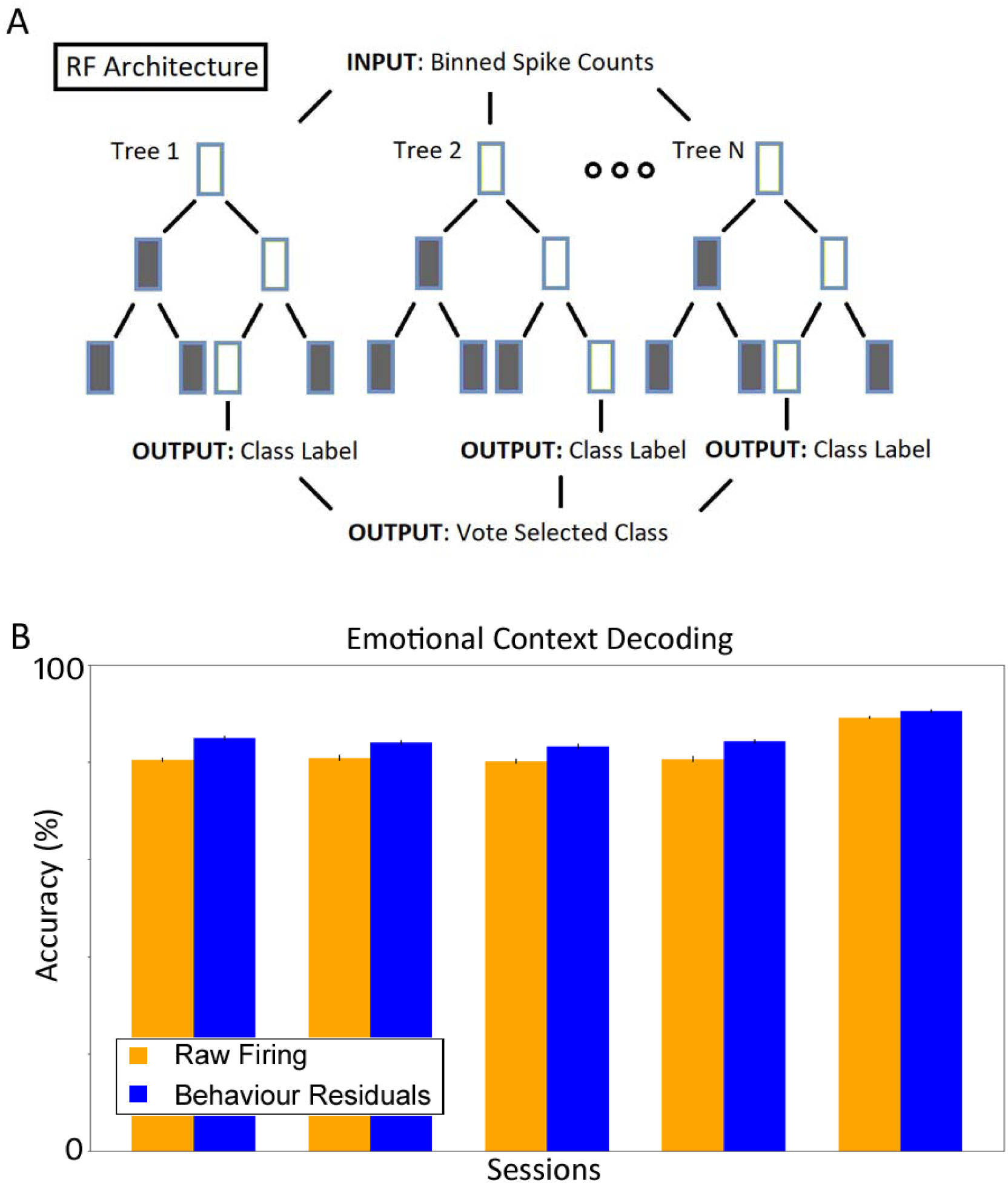
ACC ensemble encoding of emotional context and the impact of context-specific behaviors. **A)** Schematic of the RF architecture: Inputs are spike counts and the output is the predicted emotional context. **B)** Emotional context decoding was performed by RFs using raw firing rates (orange bars) or using the ‘behaviour residuals’ (blue bars) which are the residual spike counts from a GLM trained on HUB-DT labelled behaviors as factors.

### Remapping of behavioral representations by emotional context

The remainder of the study was focused on quantifying the shift in behavioral representations across emotional contexts. To provide an overall impression of how ensembles represented the same behavior across contexts, the ensemble activity was reduced to 3 dimensions for visualization purposes. Each dot in the space represents the ensemble activity pattern during a single time bin with the colour of the dot denoting the emotional context from which it was derived (Figure 9). The time bins associated with the 3 most common behaviors across the 3 contexts (large dots in the left panels) were coloured based on the context in which they occurred. If the ACC represented behaviors independently of the context, the large dots (behaviours) would have formed coherent, isolated clusters in the neural space. This was not the case, as the large dots (behaviours) were always embedded within a cluster of smaller dots of the same colour. This meant that even though the behaviors were defined by statistical consistencies in limb kinematics (via HUB-DT), the way these behaviors were represented by the ACC depended on the emotional context in which they were performed. However, this does not mean that every neuron changed how it encoded a behavior. The right panels in Figure 9 plot the average spike counts during all the time bins in which the behavior occurred in each context. Note that the effect of context on behavioral representations varied across neurons, with some neurons exhibiting a large change while in others, there was no change at all. In this plotted example, the largest shift occurred when going from the F to S context relative to the S to N context, suggesting there may be differences in the degree of remapping for specific contexts.

**Figure 9:**
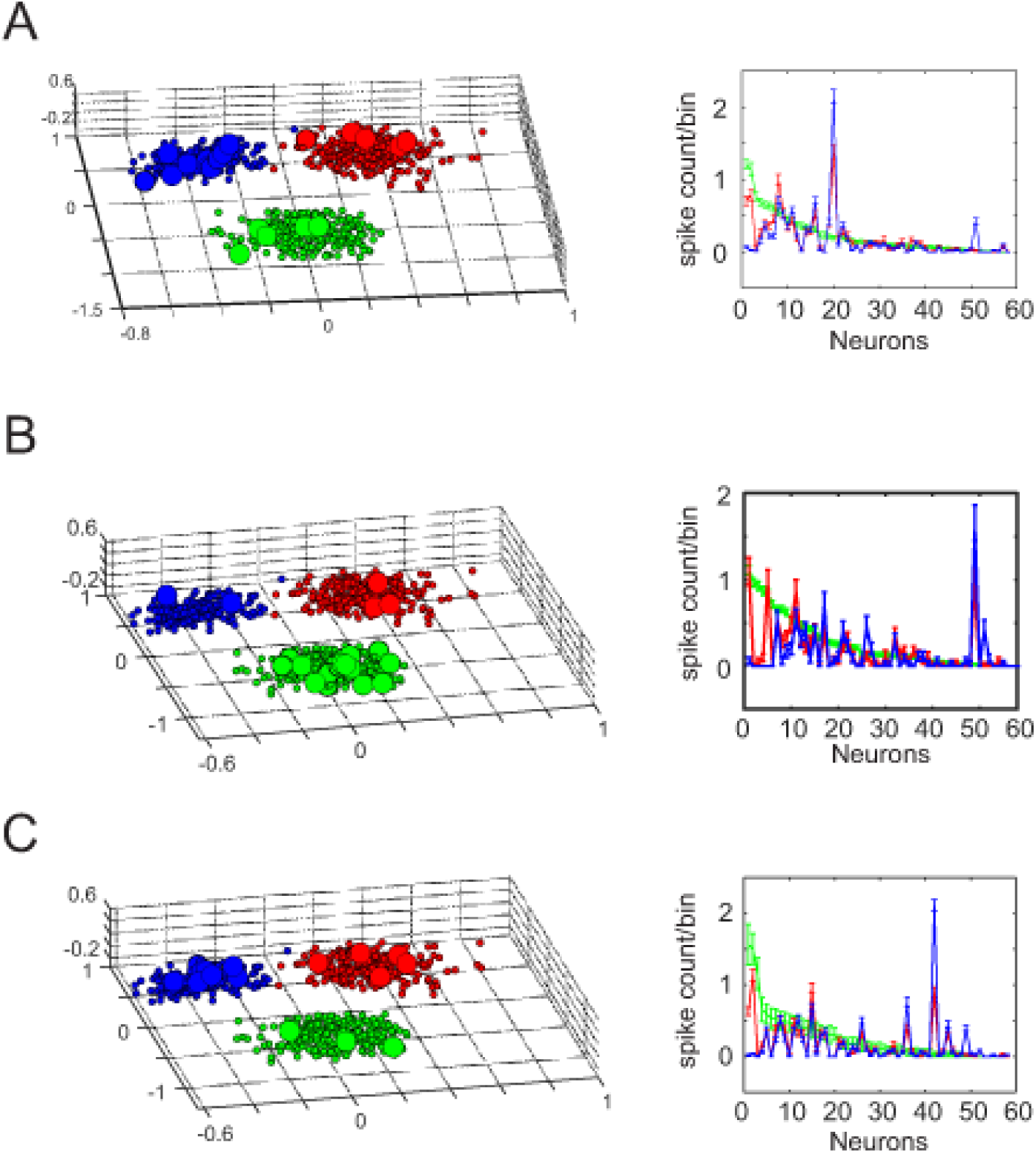
Visualizing shifts in behavioral representations across emotional contexts. A-C (Left panels) The spike counts of each neuron during all time bins from a single session were plotted on separate axes and the space was collapsed to 3 dimensions using Maximally-Collapsing Metric Learning (MCML). The large dots represent the time bins when a HUB-DT detected-behavior occurred, whereas the small dots represent spike counts in all other bins. For visualization purposes, the total number of dots was reduced by down-sampling from 0.2s bins to 5s bins via averaging. (Right panels) Each point represents the average (and s.e.m.) spike count across all 0.2s bins in which the behavior was detected in a given context, for each neuron. In this session, the order of the contexts was F-S-N and the neurons were sorted according to the magnitude of their responses in the F-context and the sorting was maintained for the remaining 2 contexts. A-C are for the 3 most frequently expressed behaviors across all 3 emotional contexts.

### Contextual Shifts in Ensemble Encoding

To quantify the magnitude of the contextual shifts, we compared the similarity of the ensemble patterns associated with a given behavior when it was performed multiple times within a single context versus when it was performed in two different contexts. To do this, the single most prevalent behavior in each session was identified and for each context, the time bins associated with that behavior were randomly split into two groups. The regularized Mahalanobis distance (DMahR) was then calculated between the two sets of points from within each context (to determine if the ensemble representation of a behavior changed within a single context) and compared to the DMahR between neuronal responses in sets of time bins associated with 2 different contexts (Figure 10). All calculations were performed in a non-reduced space that included all neurons. However, in order to compare data across sessions, a set of 40 neurons were chosen randomly for each of 100 bootstraps/session and the results were averaged. As shown in Figure 10, the distances between the ensemble patterns were much larger when the behavior was performed in two different contexts versus when it was performed within a single context (F(1,4)=43.5, p=0.003). When only the between-context comparisons were considered, the distance between comparisons involving N vs S or F vs S contexts were larger than comparisons involving the N vs F contexts (post-hoc t-tests, p<0.01).

**Figure 10:**
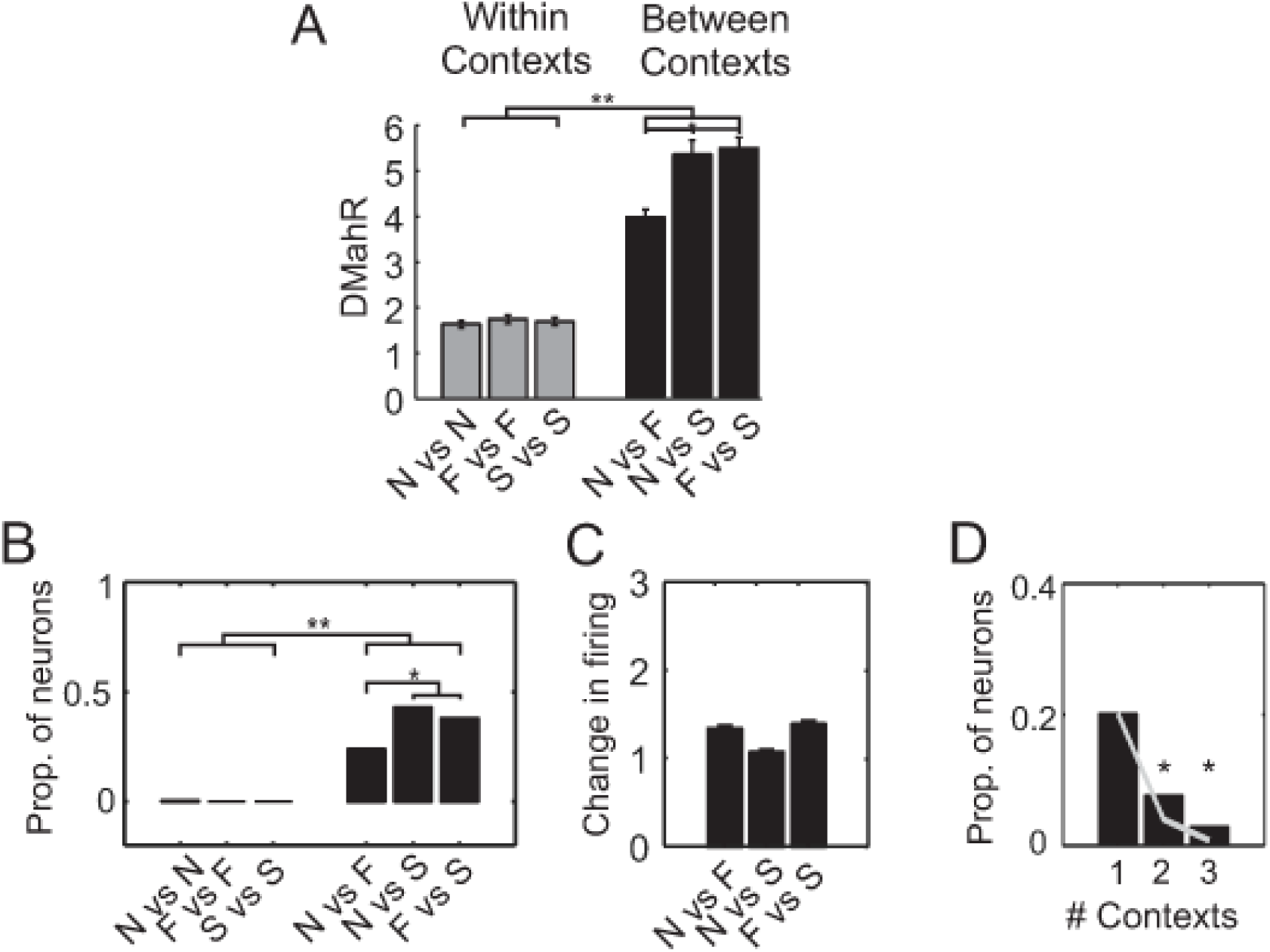
Quantifying the magnitude of change in behavioral representations across emotional contexts. **A)** Quantification of changes at the ensemble level. The most commonly performed behaviours across the 3 contexts were selected, and the DMahR was computed between spike counts associated with each behavior when the time bins were derived from within a single context versus from 2 different contexts. All cross-context distances were greater than all within-context distances (**, p<0.01) while the greatest cross-context distances were those produced by transferring into the S-context (*, p<0.05). **B)** Quantification of the proportion of individual neurons that changed responses to a behavior. The 10 most commonly performed behaviours were selected, and for each neuron a t-test was used to compare activity when the behavior was performed within a single context or in 2 different contexts. The proportion of neurons that changed how they responded to at least one of behaviors in two different contexts far exceeded the proportions that changed how they responded to at least one of behaviors within a single context (**, p<0.001). The largest proportion changed when transitioning from the N to S-contexts and this proportion was significantly larger than the proportion that changed when transitioning from the N to F-contexts (*, p<0.05). **C)** For the sets of neurons shown in (B), the average (and s.e.m.) change in spike counts associated with the behavior when it was performed in two different contexts. For perspective, the overall average spike count/bin across all neurons in the dataset was 0.34. **D)** The proportion of neurons responsive to a behavior within one, two or three contexts regardless of the type of context. The gray line gives the predicted probability of finding the observed proportion of neurons responsive in 2 or 3 contexts, given the probability of responsiveness in one context was 0.2. The stars denote significance of observed versus chance proportions according to χ ^2^ testing (p<0.001).

### The dynamics of changing behavioural representations

The ensemble distance measures gave an indication of how much the behavioral representations shifted but said little about the neuronal dynamics causing the shift. To address the question of how many neurons changed and by how much, we extracted the 10 most common behaviors in each session and calculated the proportions of neurons that responded differently to each of the behaviors within a context versus across contexts using a series of paired t-tests. Figure 10 plots the proportions of neurons that responded differently to at least one of the behaviors for the comparisons shown. In terms of the within-context comparisons, only a single neuron responded to at least one of the behaviors differently within a single context. In marked contrast, 24% (n=63), 43% (n=113) and 38% (n=100) responded differently across the N versus F, N versus S and F versus S contexts, respectively. Chi^2^ testing revealed that the proportions of cases where behavioral responses differed was always significantly greater for cross-context comparisons than within-context comparisons (all p<0.001). For the cross-context comparisons in particular, there were significantly more neurons where responses to a given behavior changed between the N versus S than N versus F contexts (χ ^2^=21, p<0.001) or the F versus S contexts (χ ^2^=12, p<0.001). Figure 10C plots the average magnitudes of the absolute change in spike counts for all cases where a neuron significantly changed its response to a behavior. Although an ANOVA revealed a highly significant overall difference (F(5,1577)=18, p<0.001), this effect was driven by the large differences between all the within-versus cross-context comparisons and post-hoc testing revealed that none of the cross-context comparisons differed significantly from each other (Figure 10C).

### Stability of Behavioural Representations

Next, we considered the opposite question of how stable single-neuron representations of behaviour were across contexts. Specifically, if a neuron was responsive to a behavior in a given context, how likely was it that it would remain responsive across 2 or all 3 contexts? In terms of this question, one drawback of the GLM and t-test approaches above was that they necessarily involved contrasts between various pairs of task epochs. Instead, we used t-tests to compare spike counts in all bins a behavior occurred within a given context to all bins it did not. Of note, this approach also accounted for the global shifts in ensemble dynamics that occur when switching from one context to another (Caracheo et al 2018). Using this approach, it was possible for a neuron to be responsive to a behavior in *any* of the 3 contexts or in *all* of the 3 contexts. The probability of the former case was 0.2, whereas the probability in the latter case was only 0.03. This meant that at least in terms of the 10 most common behaviors present across all 3 contexts, nearly 10x as many ACC neurons were responsive to a behavior in a contextually dependent manner than in a consistent manner. This raises the question of whether the probabilities were higher than chance. As the average probability that a neuron was responsive to any of the 10 behaviors in a single context was 0.2 (Figure 10D), one would expect that the probability of being responsive in 2 contexts by chance should be 0.2^2^ (0.04) and in 3 contexts, 0.2^3^ (0.008). The observed probabilities for neuron being behaviorally responsive in 2 (observed=0.08, expected=0.04; *χ ^2^*=31, p<0.001) or 3 (observed=0.03, expected=0.008; *χ ^2^*=30, p<0.001) contexts while very low, were nevertheless significantly higher than chance (Figure 10D). In other words, the neurons tended to be slightly more consistent/stable than would be expected if behaviorally responsivity was allocated in a completely random manner in each subsequent context.

### Behavioural ‘Syllables’ and ‘Words’

Behaviours do not occur in isolation, but rather are linked together in groups over time. Hence, a potential reason for the large contextual shifts in the behavioral responses could be that the behaviors were linked together in different ways in each context and the neurons were sensitive to the overall sequence of behaviors. One way to conceptualize this would be to consider the discovered behaviors as ‘syllables’ that were linked together into longer sequences that formed unique behavioral ‘words’ (e.g. Wiltschko et al 2020). In order to determine if the behavioral words were consistent within and across contexts, we again selected the 10 most common behaviors and plotted them along with behaviors that immediately flanked them. Figure 11A shows that the behaviours (colours) preceding/following a target behavior tended to be similar across all occurrences of that behaviour both within and across contexts. In order to quantify this effect, matrices of sequence labels like those shown in Figure 11A were constructed for each session and correlations between all occurrences of a behavioural word both within and across contexts were calculated. An identical series of correlations were performed on sets of randomly generated behavioral word matrices. The average absolute correlations between multiple instances of behavioural words were significantly larger than the correlations between randomly generated behavioral word matrices of the same dimensions (F(11,599)=62, p<0.0001, post-hoc multiple comparisons, p<0.01). This demonstrates a surprising degree of consistency in the sequences of behaviours emitted throughout the sessions. By extension, this rules out inconsistencies in behavioral sequences as a potential cause of the instability of behavioural representations by ACC neurons.

**Figure 11:**
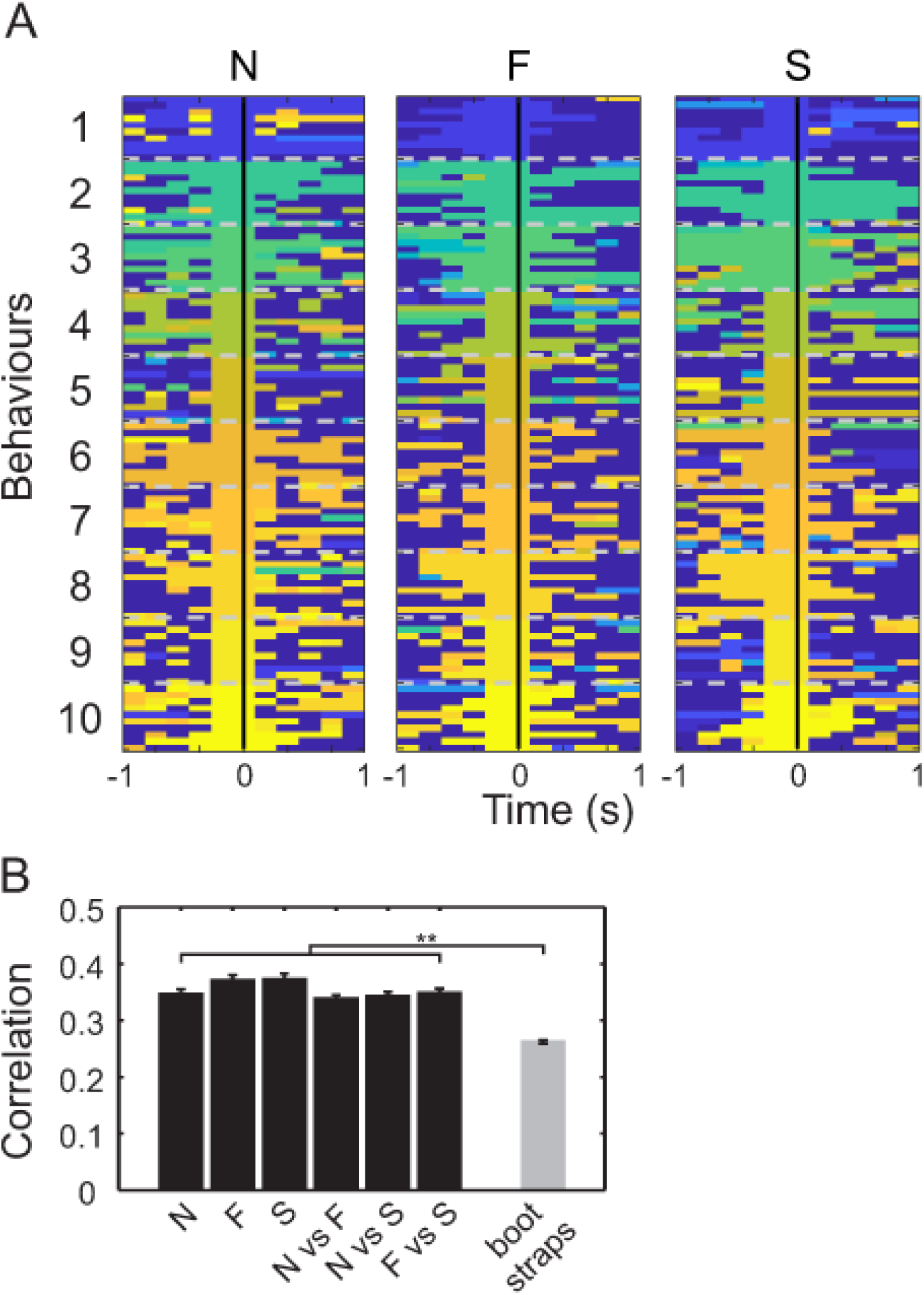
Linking behaviors through time. **A)** For a single session, 10 randomly chosen occurrences of the top 10 most common behaviors were extracted along with the behaviors in the 5 x 0.2s bins flanking each respective occurrence. Every behavior was then assigned a different colour. These colours were only used to differentiate behaviors and conveyed no quantitative information about a behavior or how similar behaviors were to each other. Each line represents one occurrence of a given behavior (centered at time 0). The horizontal gray dotted lines separate the 10 occurrences of one behavior from the 10 occurrences of another behavior. The sorting of the 10 behaviors were maintained across the 3 contexts, although the specific occurrences necessarily varied between contexts (N=Neutral context; F=Food context, S=Shock context). Since the colours were relatively similar across all occurrences of a given behavior, it meant that the sequence of behaviors surrounding the target behavior were consistent throughout the session. When viewing this plot, it is important to note that behaviors differ in adjacent 0.2s bins even though every behavior was defined by limb movements as slow as 0.5Hz; Behavioural labels are instantaneous labels of variable length processes, and each bin is assigned the most probable behaviour as a label. **B)** Average correlations between occurrences of behavioral words within and across pairs of contexts (black bars). Average correlations between randomly generated behavioral word matrices (bootstraps) of identical dimensions (gray bars).

## Discussion

In this study we developed an automated behavioral discovery pipeline that, when linked to electrophysiology data, enabled us to perform comprehensive correlation analysis of how ACC neurons flexibly encode behaviors. It was possible to at least partially disambiguate emotional context signals from behaviors and we found that the tracking of context-specific behaviors could not account for the separation of emotional contexts by ACC ensembles. However, the unique emotional states induced in each emotional context had a strong impact on the way the precisely-defined, naturally occurring behaviors were represented. The results imply that ACC representations of behavior are intimately tied to, or contextualised by the emotional state experienced when the behavior is performed.

### Emotional Regulation

There are a number of reasons to believe that the ACC is involved in the emotional regulation of behavior. On one hand, there is an extensive literature documenting the role of the ACC in emotion. This includes functional imaging studies in humans that have provided clear evidence that the ACC differentially encodes emotional states (Shackman et al 2011) and that dysfunction of the ACC contributes to a number of emotional disorders (Jaworska et al 2015; Drevets et al 2008). At a cellular level, ACC neurons encode internal states such as stress, fear, pain and hunger (Padilla-Coreano et al 2016; Quinn et al 2008; Guiustino et al 2015; Johansen et al 2001). This is consistent with anatomical data showing that the ACC has extensive interconnections with a widespread network of regions that generate and control emotional and autonomic states, including the amygdala, midbrain, brain stem and spinal cord (Hoover et al 2007; Gabbott et al 2020; Burgos-Robles et al 2017). In rodents, disruption of these pathways alters the expression of emotionally driven behaviors (Jhang et al 2018; van Heukelum et al 2021) while in primates ACC lesions result in reduced reactivity to threatening stimuli and reduced behavioral output (Rudebeck et al 2006; Bliss-Moreau et al 2021). In humans naturally occurring damage to the ACC can lead to the emergence of akinetic mutism (Devinsky et al 1995) whereas more delimited, precise surgical removal of the ACC and surrounding cortex (anterior cingulotomy) has proven to be beneficial for reducing extreme emotional reactions to severe types of pain or to anxiety-proving stimuli (McGovern et al 2017; Brotis et al 2009).

In addition, there is a largely separate literature highlighting a role for the ACC in the control of behavior. This includes anatomical studies documenting the connections between the ACC and regions that generate behavior, such as the motor cortex, thalamus, striatum and spinal cord (Leow et al 2022; Takeuchi et al 2022; Brockett et al 2020). Likewise, electrophysiological studies have shown that ACC neurons respond to the movements of individual limbs as well as a wide range of actions and behaviors (Jung et al 1998; Seo et al 2012; Lindsay et al 2018). These behavioral correlates can be modulated by a variety of factors, including the environmental context, task context as well as the effort involved in executing the behavior and the outcome it yields (Sachuriga et al 2021; Porter et al 2020; Bryden et al 2019; Caracheo et al 2018; Ma et al 2016). The confluence of emotional and behavioral information within the ACC supports the hypothesis that this region plays a central role in the emotional modulation of behavioral output (Seamans & Floresco 2022). The present study attempted to illuminate how ACC neurons combined information about emotional states and behaviors.

### Defining and Extracting Behaviour

Behaviour can be a somewhat general and ambiguous term. Here we used a specific operational definition, which was changes in pose (i.e. relative body part locations) that evolved over a range of timescales. Our automated behavioral discovery tool, HUB-DT, could reliably categorize and label statistically separable behaviours. We referred to the process of unsupervised categorisation as behavioural ‘discovery’ because HUB-DT clustered a range of behaviours, some of which were not readily differentiable to human observers simply from observation of video. Indeed, it was often subtle variations in one or two limbs that separated context specific behaviours from visually similar behaviours present in other contexts. The application of HUB-DT provided what could reasonably be considered the full spectrum of observed behaviours on this task.

Our operational definition of behaviour differs from the approach of defining behaviour according to the animal’s goals. While some of the behaviors detected by HUB-DT were goal-directed, for others the relationship was less obvious. This was partly due to the fact that the 3-valence task involved Pavlovian contingencies that did not depend on the rat’s behavior. As a result, the only truly goal-directed behavior was pellet retrieval. The other challenge was that because naturalistic behaviors flowed continuously from one to another, decisions about behavioral segregation were difficult. For this reason, it may be more appropriate to consider the behaviors detected by HUB-DT as sets of movement ‘syllables’, similar to the approach taken in correlational studies of motor output (Lashley 1951; Bizzi et al 2013). These sequences of behavioral syllables could be combined into unique behavioural ‘words’ that would be perhaps more similar to what a human observer would consider as distinct behaviors. We found that on the 3-valence task, rats often performed the same behavioral syllables in a consistent pattern throughout the session. Within these patterns, the same behavioral syllable was sometimes repeated at a high frequency, while in other instances the behavioral syllables transitioned in adjacent 0.2s bins. It is important to note that although HUB-DT could theoretically endow each 0.2ms bin with a unique behavioral label, it did not define behaviors as transitions between poses in adjacent 0.2s time bins. Instead, each instance of a given behavioral syllable corresponded to a single point within a specific cluster that was defined by pose projections across a range of frequencies from 0.2s to 2s, and is therefore informed about movement beyond the bounds of that single bin. Behavioral syllables could transition in adjacent 0.2s bins because identified behaviors neither dwelled nor transitioned at fixed, well-defined intervals. For some perspective on this issue, consider the example of an animal running.

Running is often interspersed with brief interruptions, stutters or even transitions to walking and the animal may transition from running to stumbling and back to running in less than a second even though a complete running stride cycle is longer than 1s.

### Representing Behaviour in ACC neurons

One way to verify if the behavioral boundaries were appropriate was to evaluate if they were associated with changes in the activity of ACC neurons. We found that there were robust responses to the wide variety of behaviours expressed on this task. For many context-specific behaviours identified by HUB-DT, there was a closer correspondence between ACC firing than those scored by a human observer. We acknowledge that this is not a strict comparison between HUB-DT and all ‘human observable’ behaviour, but rather to a set of general, easily observed behaviours not tailored to any specific task. We have already demonstrated that ACC neurons track changes in limb positions (Lindsay et al 2018), but this factor alone could not fully account for our results as the average relationship between firing and behavior (i.e. r^2^) was stronger than between firing and limb positions shown in Lindsay et al (2018). It is of course still possible that ACC neurons were combining representations of limb positions into more complex representations of behaviors.

### Emotional Valence and the Modulation of Behaviour

Consistent with our prior study (Caracheo et al 2018), distinct ACC ensemble state patterns were associated with each emotional context of the 3-valence task. Each context-specific pattern was initiated by the first tone within a block and remained in place throughout every subsequent time bin, including the ∼50s inter-stimulus intervals when no tones or outcomes were presented. Although HUB-DT detected context-specific behaviors, these contextual representations did not arise because the neurons were tracking these behaviors. We can say this because the analyses involving residual firing matrices showed that emotional context selectivity was largely unaffected by removal of behavioural correlates. Therefore, the context-specific patterns were likely representing the emotional reactions to the unique outcomes themselves (Caracheo et al 2018).

ACC ensembles produced clear and separable representations of each emotional context, even though individual neurons were sensitive to tones and outcomes of more than a single valence. This conclusion is in seeming contrast to the findings of Monosov et al, 2017, who identified sets of valence-specific ACC neurons in primates during a two-valence task. However, the discrepancies may come down to differences in the operational definition of neuronal responsiveness and the specifics of the analysis used. In Monosov et al., the definition of responsive neurons was limited to monodirectional modulation of a neuron to outcomes of a single valence. Neurons which responded in more complicated ways, or responded to multiple task factors were not accounted for, or were categorized as responsive to only a single one of these factors. In addition, their analyses did not take into account possible transformations of the responses similar to what we observed here. Nevertheless, both studies provide clear evidence that ACC neurons are informative of valence in addition to other stimuli and task factors.

### Mixed Selectivity and Population Encoding

The observation that behavioral representations were intermixed with emotional context representations of multiple valences is consistent with the concept of ‘mixed selectivity’. Mixed selectivity is now well-established in cortical encoding analysis and describes how multiple task factors can be represented by individual neurons and distributed across a neuronal population (Stringer et al 2019; Kira et al 2023). These structured representations are consistent and robust, but are comprised of variable groups of neurons with intermittent participation. A growing body of work including some recent advances in neural network design (Rigotti et al, 2013, Barak et al., 2013, Fusi et al, 2016, Nogueira et al, 2023, Mazzucato et al, 2016; Koay et al 2022) suggests that networks of neurons possessing mixed selectivity offer advantages in terms of flexibility and stability. In particular, clustering multiple representations within a given region of neural space affords networks the ability to construct more detailed and nuanced representations of complex task situations (Musall et al 2019; Bernardi et al 2020; Koay et al 2022) and improves network capacity. Within this framework, neurons participate in many representations, and switching between them is a process of reweighting the responses across the population. As a result, the pairings between represented stimuli and individual neurons are not necessarily linear, nor do they vary smoothly. In ensembles of neurons with mixed selectivity, consistency is expressed in terms of cluster distances and boundaries in neural space (i.e. the representational geometry). Further progress in describing these ensemble representations may therefore be possible by implementing approaches such as direct geometric analyses (Driscoll et al 2017; Gallego et al, 2020, Williams et al, 2021, Nogueira et al, 2023, Qi et al, 2017) and dynamical systems analysis (Linderman et al., 2016, Pandarinath et al, 2018).

In the present study, the effects of mixed selectivity were evident in multiple analyses. Perhaps most obviously, it accounted for the overlapping response distributions. Although each neuron tended to encode multiple behaviors, even the strongest behavioral correlates accounted for only a relatively small portion of the variance in firing. Furthermore, the responsivity of neurons to a given behavior was not an all-or-none phenomenon but was smoothly distributed across the population (Figure 7). These properties create a scenario where individual neurons weakly encode multiple task events and changes in the emotional context shift the relative strengths of each representation. In the capacity to convey information, the functional unit is the ensemble, rather than the individual neuron, and as an ensemble, the representation of behaviour, emotional context, and other task factors is highly consistent and robust.

### ‘Context’ Encoding as a General Principle

Contextual shifts in ACC representations are not limited to ‘emotional contexts’ as changes in the task context (e.g. a task rule such as the order of actions or direction of travel) or the physical context (i.e. environment) also change how actions are represented by ACC neurons. Ma et al (2014; 2016) examined the mechanistic basis of contextually-induced remapping by studying how neuronal responses to 3 operant actions varied when the actions were performed in a different order (i.e. task context) or different enclosure (i.e. physical context). The remapping was structured in such a way that all 3 action clusters were shifted in the neural ensemble space by the same amount and the relative geometry of the 3 action clusters was preserved. This was possible because there appeared to be a fixed amount of ‘responsivity’ for the actions that was distributed across the population in each context such that when the context changed, the total amount of responsivity remained constant but it was redistributed across the population (Ma et al., 2014, Ma et al., 2016). All neurons seemed to encode all events to varying degrees, but not all neurons were affected by changes in context in the same way (Figure 9). In other studies, we found that the extent and the magnitude of the shifts were dependent on how the context change was experienced. When a change in context was immediate and obvious to the animal (e.g. when the animal was moved between different physical environments), the changes in the responses of single neurons tended to be highly synchronized across the population (Hyman et al 2013), whereas in cases where the animal slowly accumulated information that the context had changed, such as during gradual trial-and-error learning (Seamans & Emberly 2020) or learned extinction (Ruso et al 2021), the changes in the responses of single neurons were much more desynchronized.

### Emotional ‘Gain’ and Contextually-appropriate Behaviour

Given the extent of contextual remapping observed in ACC, it seems unlikely that the region is dedicated to the processing of a specific type of information. Instead, we have argued that the ACC may imbue events, behaviors or sensations with a type of emotional, or contextual ‘gain’ (Floresco & Seamans 2022). Such a view is consistent with the effects of ACC lesions in primates that cause reduced reactivity to threatening stimuli and reduced behavioral output, as well as with the emergence of akinetic mutism in patients with severe ACC damage (Devinsky et al 1995; Rudebeck et al 2006; Bliss-Moreau et al 2021). To be clear, we are not saying that the ACC directly produces behaviors or emotions. Rather, it exerts a biasing influence on which stimuli or internal-states modulate behavior and how vigorously or persistently the behavior should be performed. The shifts in behavioral responses in the 3 contexts could be examples of how this process is instantiated at a neuronal level. The emotional modulation of behavioral representations may be the mechanism used by the ACC to inform downstream regions about the types and intensities of behaviors that should be emitted in a given context. This process ensures that behavioral output is appropriate to attain a goal, which is as relevant for survival in naturalistic settings as it is for success in social situations.

## Methods

### Experimental Subjects

Four male Long-Evans rats (Charles River Laboratories, Montreal) weighing between 400 and 470 g were used. They were housed in an inverted 12-h day/12-h night cycle and were food restricted to 90% of their free-feeding weight but given unlimited access to water. Tetrode array implant surgery was performed before any behavioral training began. All procedures were conducted in accordance with the Canadian Council of Animal Care and approved by the Animal Care Committee of the University of British Columbia.

### Behavior

Recording and training sessions took place inside a custom-made behavioral chamber (30 cm x 25 cm x 60 cm) with a pellet dispenser that dispensed 45 mg sweet dustless precision pellets (BioServ) into a magazine cup, a shock grid and a tone emitting speaker that were all connected to a Med Associates (St. Albans, VT) control box and to a PC workstation running Med-PC software (St. Albans, VT). Before training began, rats were habituated to the chamber and accustomed to eating food from the magazine cup. They were also exposed to foot shocks in order to determine the intensity of the shocks for the experiments. The shock duration was 500 ms and the intensity was varied between 0.5 - 0.7 mA and was chosen based on the minimum level required to elicit a noticeable behavioral response. After rats had initial exposure to the chamber and to the two outcomes, tone conditioning began. Three distinct frequency (4,6,8 kHz) tones were assigned to three possible outcomes: food, shock or nothing.

Prior to conditioning, the tones evoked no noticeable behavioral reactions. Tones were randomly assigned to an outcome for each rat, but once conditioning began, tone-outcome contingencies remained unchanged for the duration of the experiments. During the initial training period, tone-food pairings were conducted for 5 days, followed by 2-3 days of tone-shock and tone – no outcome pairings. Rats were exposed to at least 3 sessions where they received all three contingencies in consecutive blocks before any of the recordings used in this study took place. A session consisted of 3 blocks with the order randomly selected from all possible combinations. Each block consisted of 30 trials where the tone was presented for 5 seconds and the outcome was delivered immediately thereafter. The average ITI was 49s (range 25-85s). The data set was constructed from 5 selected recording sessions from 4 animals with varied block orders. The animal’s behavior in each block was video recorded using a dual camera set up that captured side and bottom-up views.

### Surgery and Data Acquisition

The electrophysiological recording data have been published in Caracheo et al (2018), using a different series of analyses. Rats were surgically implanted with custom built 16 tetrode hyperdrive arrays. They were anesthetized under iso-flurane gas, their skull was surgically exposed and a 4 mm x 3 mm hole was drilled. The dura was removed to expose the brain around coordinates +3.0 mm from Bregma and ±0.5 mm from the midline. The implant was positioned over the area and fixed to the skull with 11 skull screws and dental acrylic. Two additional screws used as ground wires were placed in the posterior skull. Tetrodes were lowered ∼1000 μm on the day of surgery and then the rats were given 1‒2 weeks of recovery. Tetrodes were advanced up to 1000 μm more to their target prior to the first recording session. In between sessions, tetrode drives were turned between 20 – 50 μm to maximize the units recorded and obtain different populations. When the experiments ended, rats were perfused and their brains collected and sliced on a cryostat. Slices were mounted on slides and viewed under a microscope to confirm the anatomical locations of tetrode tracts. Based on tetrode advancement records, the positions were estimated to have been mostly in the medial wall, within the ACC up to the border of the prelimbic cortex (PL).

Tetrodes were attached to EIB-36TT boards, plugged into two HS-36 headstages and connected via tether cables to a Digital Lynx 64-channel system and then to a PC workstation. Electrophysiological data and behavioral events were captured using Cheetah 5.0. Files were exported into Offline Sorter (Plexon, Inc.) and were manually sorted based on three-dimensional projections of wave form peaks, valleys and principal components. Once cells had been sorted, they were exported to Neuroexplorer 4 (Nex Technologies) and then to Matlab or Python for further analysis.

### Video Data Collection

Subjects were recorded by a custom setup of two synchronized colour video cameras, one camera side-on to the enclosure and another camera below. Automated feature tracking was performed on combined videos using DeepLabCut (Mathis et al., 2018). The resulting two sets (one for each camera angle) of 2-D tracking were combined into a single 3-D tracking. The central point of the skeleton was set to the centre point of the rat’s body. The x-y-z location of this point gave the spatial location of the animal at each time bin. The positions of 9 body parts (head, mid-shoulder, front left limb, front right limb, pelvis, rear left limb, rear right limb, base of the tail, mid-point of tail) were calculated relative to the centre point of the body. The frame rate of the video (5Hz) and the neural data (200ms bins) were synchronized. The alignment was set by paired pulses to the video and electrophysiology recording rigs, matching the timing to within 1ms.

A set of human labelled behaviours was generated by manual annotation. These annotated behaviours were a set of standard behaviours scored from video. Behavior in each time bin was categorized as belong to one or more of the following; locomotion, rearing, grooming, foraging/feeding, sniffing (i.e. interacting with the environment through head, whisker movements), and resting. A human observer fully annotated 2 of the 5 total sessions used in this analysis. The human observer-generated labels were propagated across the remaining sessions by a trained Random Forest (see below for details). Since one of the human-annotated behaviours, feeding, was by definition context specific, it was omitted from context purity comparisons.

### Hierarchical Unsupervised Behavioural Discovery (HUB-DT)

3-d tracking data from DeepLabCut was post-processed by subtracting out the location of the animals’ centre of mass. This resulted in a set of tracking data where all body parts had coordinates relative to the animals’ centre of mass. The data was agnostic about location in the enclosure as well as the direction or orientation of the animal in the enclosure. HUB-DT quantified behaviors based on an approach similar to that used in Motion Mapper (Berman et al., 2016). Specifically, a spectrographic projection via Morlet wavelets was used to discretize spatial and temporal features together into an expanded behavioural space. This involved first performing a Fast Fourier Transform on the xyz coordinates of each of the nine body parts and then separately convolving these with 5 logarithmically distributed frequency Morlets (ranging from 5Hz to 0.5Hz). This resulted in a full behavioural space of 9x3x5 (135) dimensions that quantized the changes in the relative positions of each body part across the 5 selected timescales.

HUB-DT then applied dimensionality reduction to the full behavioural space as an aid to visualization and as preparation for clustering. t-SNE (t-distributed stochastic neighbourhood estimation, Van der Maaten et al., 2014) was used in order to maximally preserve local structure in the reduced space. As an improvement on vanilla t-SNE, we used the openTSNE python package that utilizes approximate neighbourhood calculations, interpolation, and multi-phase optimisation to greatly improve performance and reduce computation resources. This resulted in the data being embedded in a reduced two-dimensional manifold, which we visualized via a density map showing the distribution of the data in the reduced space. This dimensionality reduction step has minimal impact on informative clusters as highlighted by Berman et al. (2014). The reduced dimensionality projection by t-SNE was used as input to the clustering algorithm, HDBSCAN. This density-based clustering algorithm was used to divide our data into a set of putative behaviours in an unsupervised way. Density-based clustering separates points via neighbourhoods of local density (calculated as reachability distance), resulting in a set of clusters without prior definition of cluster shape or number. The HDBSCAN package (McInnes et al 2017) modifies canonical DBSCAN by constructing a minimum-spanning tree of reachability distances to assign points to clusters of variable density via a hierarchy of density values. This hierarchy was used to produce cluster assignments of variable depth in the spanning tree, resulting in behavioural discriminations at different levels of granularity.

### Behavioral word analyses

For each session, the 10 most frequently expressed behaviors detected by HUB-DT across all 3 contexts were identified (i.e. the target behaviors). The behavioral labels in the 5 x 0.2s bins preceding and the 5 x 0.2s bins following each instance of a target behavior were then extracted. When the center bin containing the target behavior was included, this yielded a vector of 11 bin labels that we referred to as a behavioral ‘word’. Ten instances of behavioral words surrounding a target behavior were selected randomly from each context, yielding a behavioral word matrix containing 330 values (11bins/word, 10 word instances/3 contexts). In total across all sessions, there were 25 unique behavioral labels that were translated into 25 unique colors by the *image* command in Matlab. To perform the correlations of behavioral words, for each behavioral word, correlations between the sets of labels across the 10 instances within each context (N,F and S) as well as across all pairs of 2 contexts (N vs F,N vs S, F vs S) were calculated, yielding 6 correlation values/behavioral word. As noted above, each behavioral word matrix had 330 values and there were 10 behavioral words chosen for each of the 5 sessions. For the bootstrap correlation analysis, matrices of identical dimensions were created using the *randperm* command in Matlab, based on 25 unique behavioral words. The identical sets of correlations were performed on these bootstrap matrices as described above for the actual behavioral word matrices. This was repeated 50x and the results averaged. However, rather than reporting the average across the 6 comparisons as for the real data, the 6 average correlation values were averaged into a single value and plotted. In all cases, the correlations were converted to absolute values. The absolute correlation values were compared using a one-way ANOVA and post-hoc multiple comparisons with critical values derived using Tukey’s honestly significant difference procedure.

### GLMs for single unit statistics

The generalized linear models (GLMs) used here treated recorded firing in each time bin as a Poisson sequence with a logarithmic link function to the factors. The GLMs were set up as multi-factor predictors, thus modelling the relationship between binned neuron firing and sets of categorical factors. These factors were one-hot encoded sets of categories which translated the 3 emotional blocks (S-context,F-context,N-context), 3 tone categories (Shock-tone, Food-tone, Neutral tone) or behavioral categories identified by HUB-DT, into a series of binary vectors. The strength of the relationship between a factor and firing in the model was measured by the r^2^ value. ‘Responsive’ neurons were selected by thresholding r^2^ values; the thresholds were set to 0.01 (i.e. 1% of firing, and 0.9 or 90% of firing based on outlier analysis). As separate GLMs were used for each factor, any overlap in the responses to multiple factors was based on post-hoc evaluation.

### Purity Measures

Context specificity was measured by a two-way cluster purity measure, which was calculated between two sets of labelled clusters. In this case, one set of cluster labels was for the behaviours, and the other was for the emotional context. The purity measure was the ratio between the proportion of a given behavioral cluster that was contained within a particular emotional context cluster versus the proportion of the emotional context cluster that was accounted for by the behavioral cluster.

### Responsiveness Ratio

The responsiveness ratio was the proportion of single neurons responsive to a given factor as measured by a fit GLM. Specifically, here, the ratio gives the proportion of responsive neurons in the population to different sets of behavioural labels.

### Random Forests

A set of Random Forests (RFs) was used to decode emotional context, and behavioural labels from ensemble firing. These forests were built and trained using the Scikit-learn python package (Pedregosa et al 2011). Forests were tuned for size and subset split parameters using ‘out-of-bag’, or OOB error. Each tree in the RF was trained on a subset, or bag, of the training data and then validated on samples from outside this subset. The average of this error across all trees in the forest is the OOB error. Forests of 100 trees provided the best average results across classification tasks in this study. For raw firing decoding, each forest took as input an N item spike count vector (N = number of neurons) of ensemble firing, and predicted emotional context, or behavioural category as output. Residual firing models instead took residual firing (as generated from our GLM regression models) as input to predict emotional context. Model performance was evaluated on cross-validated folds to control for evolving patterns in the data. Reported results for all of the RF models were average performance across all folds. A similar approach was taken for training RF models to propagate manual behavioural annotations across sessions. Here the RF models took points in the full behavioural space as input, which were 9x3x5 (features by xyz by wavelet frequency) item vectors, and learned the associated manual behaviour labels as output.

### Malhalanobis Distance analysis

For each session the most frequent behavior detected by HUB-DT was identified. Then a series of bootstraps were created. For each bootstrap, 40 neurons were sampled at random and 2 sets of 3 instances of the behavior (i.e. the bin in which the behavior occurred) from each context were sampled at random (using *randperm* in Matlab), yielding 6 matrices of 3 x40 spike count bins. The regularized Mahalanobis distances (DMahR) were calculated between all the pairs of sets of matrices derived from within each context (within context DMahR). The DMahR was also calculated on a randomly chosen sets of matrices from 2 different contexts (across context DMahR). This process was repeated 100x per session. The mean DMahR from all within- and between-context bootstraps was calculated for each session and the results compared using a repeated measures ANOVA and post-hoc multiple comparisons using Tukey’s honestly significant difference procedure.

### Single neuron behavioral responsivity

The first analysis compared neuronal responses to the same behavior across pairs of task epochs. For each session, the 10 most frequently expressed behaviors detected by HUB-DT across all 3 contexts were identified and the occurrences of a behavior in each context were randomly split into two sets. Then for each neuron, paired-sample t-tests compared the spike counts associated with the two sets of occurrences within each context and also compared one of the sets chosen at random from every pair of different contexts. The p-value was set to 0.05/square root of (263*6 * 10) since there were 263 neurons, 6 comparisons for 10 behaviors. The proportions of neurons deemed to be significantly responsive to a behavior in each condition were then compared using an ANOVA and post-hoc multiple comparisons using Tukey’s honestly significant difference procedure. The second analysis compared neuronal responses to a given behavior during all bins when the behavior was performed in a context versus all bins when it was not. The p-value was set to the same value as above and the proportions of neurons responsive to a behavior within one, two or three contexts was calculated irrespective of context type. These proportions were compared using a Chi2 test of independence.

## Acknowledgements

This work was supported by funding from CIHR (MOP-137045 and PJT-159796) and NSERC (05979)

## Code Availability

Links to Github Repositories for code used in this study are linked below.

HUB-DT: https://github.com/Loken85/HUB_DT

Other Analysis: https://github.com/Loken85/ephys_ML_repo

